# Conceptual associations generate sensory predictions

**DOI:** 10.1101/2022.09.02.506283

**Authors:** Chuyao Yan, Floris P. de Lange, David Richter

## Abstract

A crucial ability of the human brain is to learn and exploit probabilistic associations between stimuli to facilitate perception and behavior by predicting future events. While studies have shown how perceptual relationships are used to predict sensory inputs, relational knowledge is often between concepts rather than percepts (e.g., we learned to associate cats with dogs, rather than specific images of cats and dogs). Here we asked if and how sensory responses to visual input may be modulated by predictions derived from conceptual associations. To this end we exposed participants to arbitrary word-word pairs (e.g., car – dog) repeatedly, creating an expectation of the second word, conditional on the occurrence of the first. In a subsequent session, we exposed participants to novel word-picture pairs, while measuring fMRI BOLD responses. All word-picture pairs were equally likely, but half of the pairs conformed to the previously formed conceptual (word-word) associations, whereas the other half violated this association. Results showed suppressed sensory responses throughout the ventral visual stream, including early visual cortex, to pictures that corresponded to the previously expected words compared to unexpected words. This suggests that the learned conceptual associations were used to generate sensory predictions that modulated processing of the picture stimuli. Moreover, these modulations were tuning-specific, selectively suppressing neural populations tuned towards the expected input. Combined, our results suggest that recently acquired conceptual priors are generalized across domains and used by the sensory brain to generate feature specific predictions, facilitating processing of expected visual input.

## Introduction

The brain is apt at exploiting statistical regularities in the world to improve the efficiency of information processing and optimize behavior (Goujon & Fagot, 2013; Gronau et al., 2008; Hunt & Aslin, 2001; Turk-Browne et al., 2005). Specifically, the brain may continuously generate predictions about future and current inputs based on previous experience. Indeed, it has been argued that the brain may implement a form of probabilistic inference (Bar, 2007; Clark, 2013; de Lange et al., 2018), combining predictions with incoming sensory information to form the most likely interpretation of the world around us. In line with this idea, neural activity in sensory areas, following visual statistical learning, appears to reflect the interplay of predictions and sensory inputs. For example, studies demonstrated attenuated neural responses to expected compared to unexpected stimuli in the ventral visual stream (Kok et al., 2012; Manahova et al., 2018; Meyer & Olson, 2011; Richter et al., 2018; for a review see: de Lange et al., 2018), often labelled ‘expectation suppression’ and resembling signatures of prediction errors as proposed by predictive processing theories (Friston, 2005; Rao & Ballard, 1999). In vision alone sensory predictions appear to be manifold, being evident for various types of stimuli and associations, such as simple features (Barne et al., 2020; Kok et al., 2017), complex objects (Manahova et al., 2018; Meyer & Olson, 2011; Richter et al., 2018), and spatial arrangements (He et al., 2022).

However, little is known whether and how predictions derived from conceptual (semantic) associations modulate sensory processing. In most prior studies (e.g., Kok et al., 2012; Meyer & Olson, 2011; Richter et al., 2018; Manahova et al., 2018) associations were learned and tested for the same visual stimuli, thus allowing for specific exemplars and low-level features to be predicted. Yet, our prior knowledge about the sensory world extends far beyond such simple regularities. We know for instance what cakes are, making them highly surprising inside shoe cabinets, without the need for exposure to a specific cake before experiencing surprise to find it in an unusual place. However, at present it is unclear whether the resulting surprise arises during perceptual processing or only at a post-perceptual level. Indeed, it is possible that such surprisal arises only after sensory processing is concluded, potentially reflected by upregulations in neural responses in anterior insula or inferior frontal gyrus, previously shown to accompany surprising input (Fazeli & Büchel, 2018; Ferrari et al., 2022; Horing & Büchel, 2022; Loued-Khenissi et al., 2020; Richter & de Lange, 2019; Weilnhammer et al., 2021). Alternatively, sensory processing itself may be modulated by conceptual priors, even though no specific exemplars or low-level features were predicted. This latter hypothesis is suggested by previous work demonstrating that (written or spoken) words prime corresponding object images (Stein & Peelen, 2015; Puri et al., 2009; Lupyan & Ward, 2013), thus implying that conceptual knowledge might be used to predict sensory input. However, object congruency (e.g., the word “cake” predicting an image of a cake) and generalizing statistical regularities via conceptual associations (knowing that cakes are usually stored in the fridge) are different affairs, with the latter involving further abstraction and predictions between different object concepts.

Here we examine whether and how priors derived from statistical regularities are automatically generalized across domains, via conceptual associations, to modulate visual processing. Moreover, we asked whether such conceptual priors, without exposure to predictable visual features, may nonetheless result in tuning specific sensory predictions. To this end, we first exposed participants to pairs of sequentially presented object words (e.g., “car” followed by “dog”). Word pairs were probabilistically associated, thus resulting in trailing words being predicted (expected) by virtue of the leading words. On the next day, participants performed a categorization task on the same object pairs, but the trailing (second) object word was replaced by an image of the corresponding object (e.g., the word “car” followed by an image of a dog). Crucially, the leading words were *not* predictive of the trailing object images. Thus, any predictions of the trailing object images must have arisen because of a generalization from the previously learned word-word transitions to the corresponding word-image pairs. Our results revealed that participants used the learned associations between word pairs to facilitate categorization of the (previously) expected trailing object images. Moreover, we found a tuning specific suppression of sensory responses, in terms of fMRI BOLD, throughout the ventral visual stream to object images which correspond to the expected words. These findings suggest that priors derived from conceptual associations result in feature specific predictions that modulate perceptual processing throughout the visual system. Thus, our results demonstrate that the predictive brain uses prior knowledge beyond concrete perceptual associations to generate and test sensory predictions, including priors derived from recently acquired abstract associations.

## Results

During a first session, participants were exposed to predictable object word pairs. In these pairs, the first word probabilistically predicted the identity of the second word, allowing participants to anticipate the trailing word based on the leading word. However, participants were not informed of the presence, nor required to learn the underlying statistical regularities. Instead, they were tasked to categorize the object pairs as congruent (both words referring to living or non-living objects) or incongruent (one word referring to a living and the other to a non-living object). On the next day, participants were exposed to word-image pairs in the MRI scanner. For each word pair, the trailing object word was replaced by an image of the corresponding object (e.g., “car” followed by a picture of a dog). Additionally, the leading words were no longer predictive of the trailing objects. Participants were asked to determine whether the trailing object image was alive or not alive. This allowed us to examine whether the learned object associations were generalized across domains from words pairs to word-image pairs.

### Word associations facilitate categorization performance of associated object images

First, we evaluated whether participants learned the word associations. Data from the learning session (word pairs) were analyzed in terms of accuracy and reaction time (RTs). Unexpected trailing words could require the same or different response compared to expected words. Thus, we analyzed unexpected words requiring the same or different responses separately, as the latter required additional response adjustments. Our results (Figure 1A) demonstrated that participants categorized the trailing words in the expected condition more accurately (96.2 ± 0.7% vs 91.1 ± 0.7%, mean ± SE; t_(34)_ = 7.798, p = 4.5e-9, d_z_ = 1.337) and faster (654 ± 5.7 ms vs 748 ± 4.7 ms; t_(34)_ = 12.426, p = 3.4e-14, d_z_ = 2.131) than the trailing words in the unexpected condition that required the same response. In addition, participants categorized the unexpected trailing words more accurately (t_(34)_ = 2.735, p = 0.009, d_z_ = 0.469) and faster (t_(34)_ = 2.845, p = 0.007, d_z_ = 0.488) when the trailing words required the same response compared to the different response. These results suggest that participants learned the associations between the word pairs and used them to predict the identity of the trailing words.

**Figure 1.**
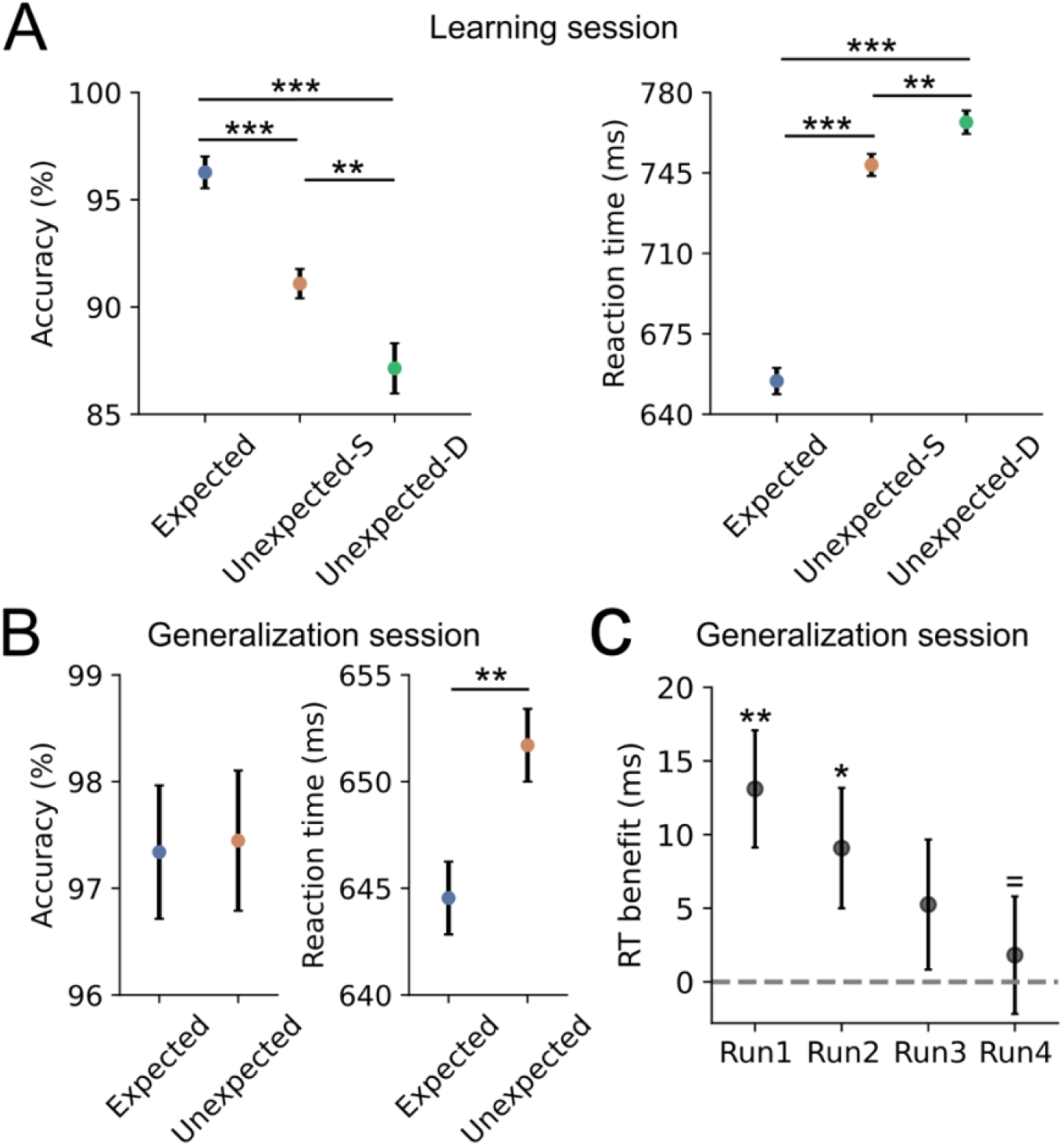
Word-word associations facilitate behavioral responses to corresponding word-image pairs. **A)** Behavioral benefits of prediction (expectation) for the word pairs indicates the learning of word associations during the learning session. Responses to expected trailing *words* were significantly more accurate (left) and faster (right) compared to unexpected words that required the same (unexpected-S) or different response (unexpected-D). **B)** Behavioral performance for the category classification task during the generalization (MRI) session. Leading words were not predictive of the trailing *images* in the generalization session. Therefore, any behavioral benefits of prediction must have been derived from the word-word associations learned during the learning session. Responses were highly accurate (left) and did not differ between expectation conditions. RTs (right) were significantly faster to expected trailing object images compared to unexpected object images. **C)** Prediction induced RT effects decreased over time until they disappeared completely by the final run. Error bars indicate within-subject SE. * p < 0.05, ** p < 0.01, *** p < 0.001, ^=^ BF_10_ < 1/3.

Next, we analyzed the behavioral data from the generalization (MRI) session to assess the spontaneous generalization of word pairs from the learning session to the word-image pairs. Overall, participants categorized the trailing images with high accuracy in both the expected (97.3 ± 0.6%, mean ± SE) and the unexpected conditions (97.4 ± 0.7%, mean ± SE). Additionally, participants also correctly rejected the no-go trials (93.0 ± 1.2%, mean ± SE), indicating good task compliance. Response accuracy for expected and unexpected trailing images did not differ significantly from each other (t_(34)_ = 0.424, p = 0.675, d_z_ = 0.073). Given the high accuracy in both conditions (>97%), this null result may reflect a ceiling effect. However, RTs for expected object images were faster than for unexpected objects (644.5 ± 1.7ms vs 651.7 ± 1.7ms, mean ± SE; t_(34)_ = 2.968, p = 0.005, d_z_ = 0.509). Crucially, while we refer to expected and unexpected object images here, the trailing objects were in fact only (un-)expected by virtue of the associations learned during the learning session, as the trailing object images were equiprobable during the generalization session. Thus, participants appeared to have generalized the associations from the word-word pairs to speed up the categorization of the corresponding expected trailing object images.

Finally, we assessed whether the prediction effect extinguishes over time, given that there were no predictive relationships during the second (fMRI) session. Thus, we analyzed the behavioral data for each run separately. Figure 1C shows that the prediction induced RT effects were evident in run 1 (13.1 ± 4.0ms, t_(34)_ = 2.828, p = 0.008, d_z_ = 0.485) and run 2 (9.1 ± 4.1ms, t_(34)_ = 2.073, p = 0.046, d_z_ = 0.356), but not in run 3 (5.3 ± 4.4ms, t_(34)_ = 1.206, p = 0.236, d_z_ = 0.207, BF_10_ = 0.458) or run 4 (1.8 ± 4.0ms, t_(34)_ = 0.480, p = 0.634, d_z_ = 0.082, BF_10_ = 0.271). Indeed, Bayesian analyses showed moderate support (BF_10_ < 1/3) for the absence of an RT effect by the last run of the experiment. Thus, the prediction induced RT effects were gradually extinguished during exposure to the word-image pairs and eventually disappear entirely, reflecting the gradual updating of predictions to reflect the non-predictive associations during MRI scanning.

### Conceptual predictions modulate sensory processing

To investigate whether neural responses to the object images were modulated by the previously learnt word pair associations, we first performed an ROI analysis targeting three distinct levels of the visual processing hierarchy: early visual cortex (EVC), lateral occipital cortex (LOC), and ventral temporal cortex (VTC); see Figure 2A. These three ROIs were defined to investigate possible prediction induced activity modulations in low-level visual areas (EVC), intermediate object-selective regions (LOC), and higher category-selective visual cortex (VTC; for details see: *Materials and Methods: ROI definition*). Our results, depicted in Figure 2B, demonstrated suppressed BOLD responses for expected compared to unexpected object images in EVC (t_(33)_ = 3.628, p = 0.001, d_z_ = 0.622), object-selective LOC (t_(33)_ = 2.782, p = 0.009, d_z_ = 0.477) and VTC (t_(33)_ = 2.851, p = 0.007, d_z_ = 0.489). To verify that our results did not depend on the specific ROI mask size, we successfully replicated all ROI results with mask sizes ranging from 50-600 voxels in steps of 50 (Figure S1). Complementary to the ROI analysis, we performed a whole-brain analysis (for details see: *Materials and Methods: Univariate data analysis)*. Figure 2C shows that expected object images resulted in attenuated brain activity throughout the ventral visual stream compared to unexpected objects. This reduction of neural activity by predictions, also known as ‘expectation suppression’, was primarily evident in the ventral visual stream, including parts of lingual and fusiform gyrus, calcarine sulcus, and cuneus. Taken together, these results suggest that the word pair associations resulted in a suppression of sensory responses to the corresponding expected object images across major parts of the ventral visual stream, including EVC.

**Figure 2.**
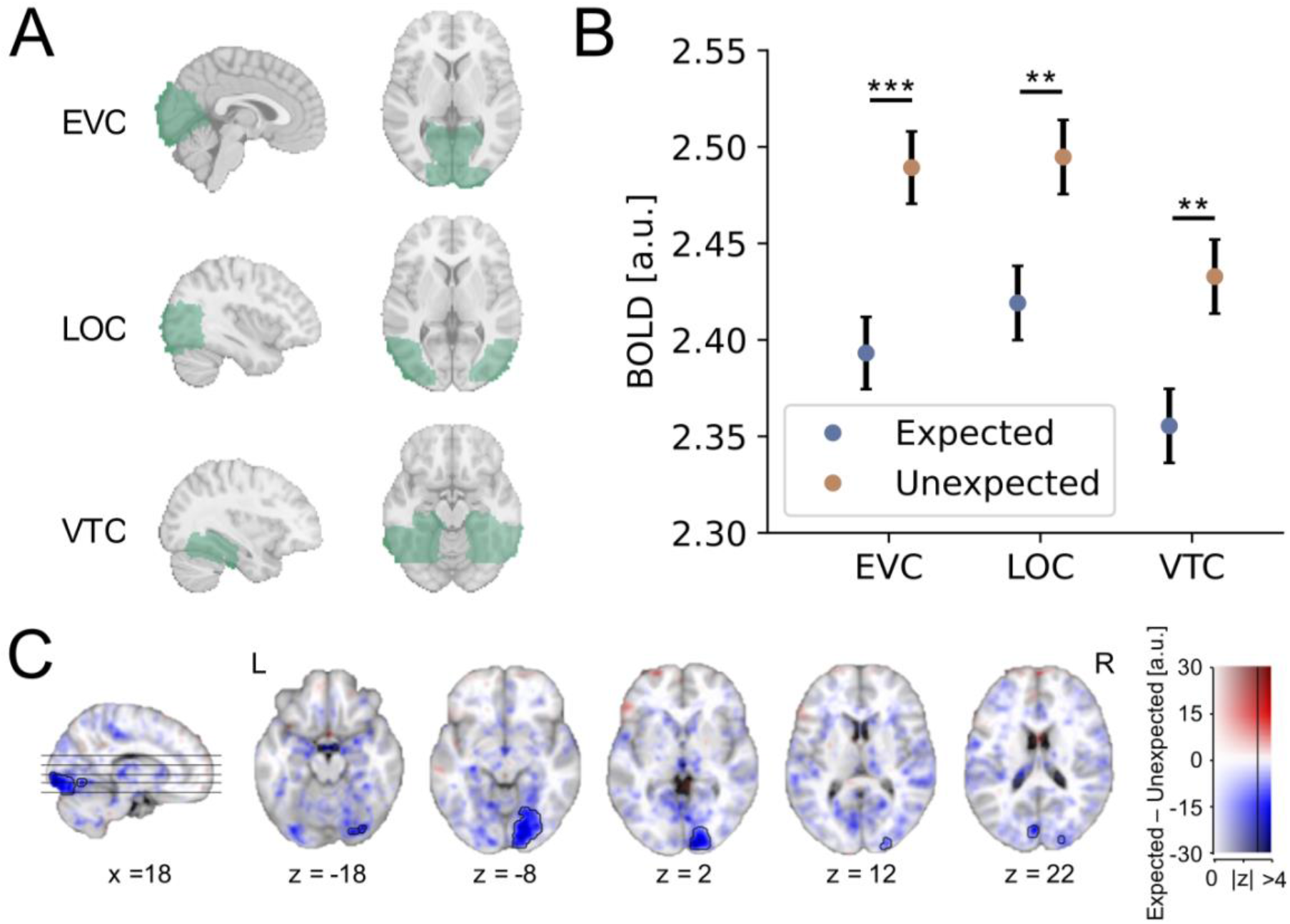
Expectation suppression across the ventral visual stream. **A)** Three anatomical masks in the ventral visual pathway: EVC (top), object-selective LOC (middle), and VTC (bottom). These anatomical masks were further constrained per participant using independent localizer data (see *Materials and Methods: ROI definition*). **B)** Averaged BOLD responses (parameter estimates) to expected (blue) and unexpected (orange) object images within EVC, LOC, and VTC. In all three ROIs, BOLD responses were significantly suppressed to the expected compared to unexpected object images. Error bars indicate within-subject SE. ** p < 0.01, *** p < 0.001. **C)** Expectation suppression revealed by whole-brain analysis. Color represents the parameter estimates for the contrast expected minus unexpected, displayed on the MNI-152template brain. Blue clusters represent decreased activity for expected compared to unexpected object images. Opacity indicates the z statistics of the contrast. Black contours outline statistically significant clusters (GRF cluster corrected). Significant clusters were observed in the ventral visual stream, including EVC, LOC and VTC.

Since behavioral prediction effects decreased and eventually disappeared by run 4 of the generalization session (Figure 1C), we also investigated the development of the neural prediction effect (expectation suppression) over time. Therefore, we performed a similar ROI analysis as above, but at the run level. Figure 3 shows an overall trend of suppressed activity for expected compared to unexpected objects within all three ROIs. Reliable expectation suppression was observed in run 2 and run 4 (all p < 0.05, for details see: Table S1), but not in run 1 and run 3 (all p > 0.215). While the run-wise estimates were noisy, given the low number of trials per datapoint, it is intriguing that no clear extinction of expectation suppression was observed during run 4 even though no behavioral facilitation was present during the same run. This suggests that expectation suppression in sensory cortex may be dissociated from the behavioral facilitation.

**Figure 3.**
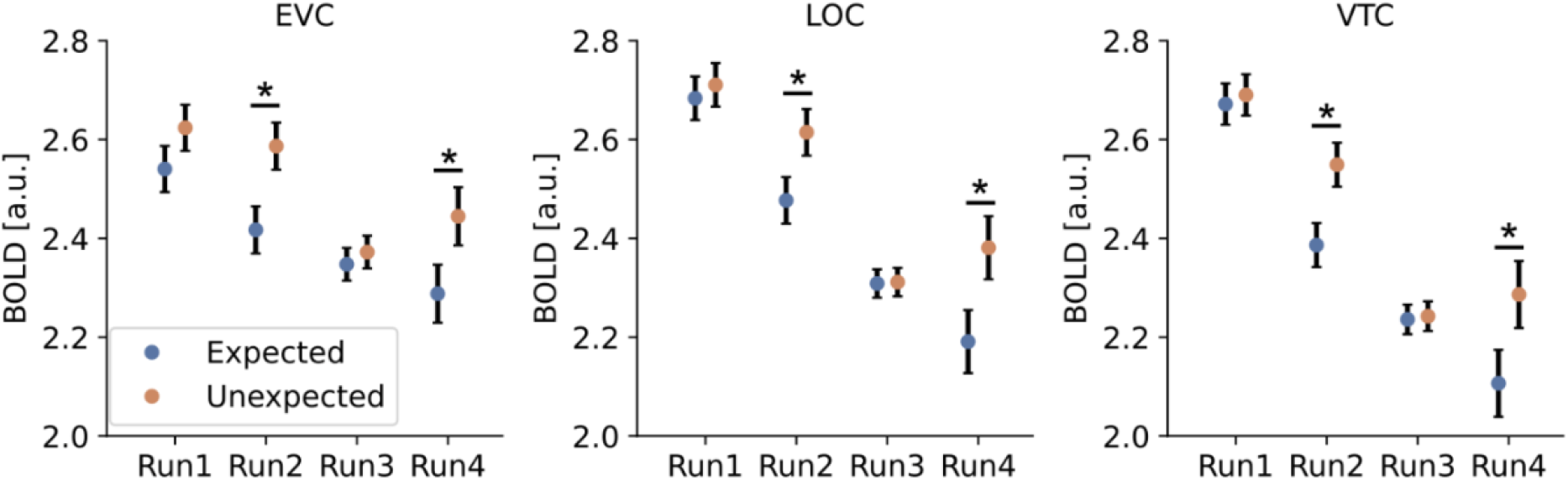
Expectation suppression across runs. Averaged BOLD responses (parameter estimates) to expected (blue) and unexpected (orange) object images for each run within EVC (left), LOC (middle), and VTC (right) across the four fMRI runs. Unlike for the behavioral effect (Figure 1C), no clear extinction of the expectation suppression effect is evident. Error bars indicate within-subject SE. * p < 0.05.

### Feature specific expectation suppression from conceptual associations

Next, we investigated whether the observed neural suppression in sensory cortex was feature specific, and thus dependent on neural tuning, or reflecting a general and unspecific upregulation of neural responses to surprising (unexpected) input, irrespective of tuning. To this end, we first generated stimulus preference ROIs by splitting voxels (neural populations) within each anatomical mask into two populations depending on whether they preferentially responded to images of living (e.g., faces and body parts) or non-living (e.g., houses and tools) objects (see *Materials and Methods: Expectation suppression selectivity analysis*). BOLD responses to expected and unexpected objects were extracted within the resulting ROIs for each category and averaged separately depending on whether the object image was of the preferred or non-preferred stimulus category. Our results, depicted in Figure 4, showed that expectation suppression was robustly present in all three visual ROIs when the category of the trailing images was preferred (EVC: t_(33)_ = 2.913, p = 0.006, d_z_ = 0.500; LOC: t_(33)_ = 4.062, p = 2.8e-4, d_z_ = 0.697; VTC: t_(33)_ = 4.710, p = 4.3e-5, d_z_ = 0.808). However, there was no reliable suppression of neural responses when the trailing images were not preferred (EVC: t_(33)_ = -0.988, p = 0.331, d_z_ = -0.169; LOC: t_(33)_ = 0.928, p = 0.360, d_z_ = 0.159; VTC: t_(33)_ = 0.992, p = 0.328, d_z_ = 0.170).

**Figure 4.**
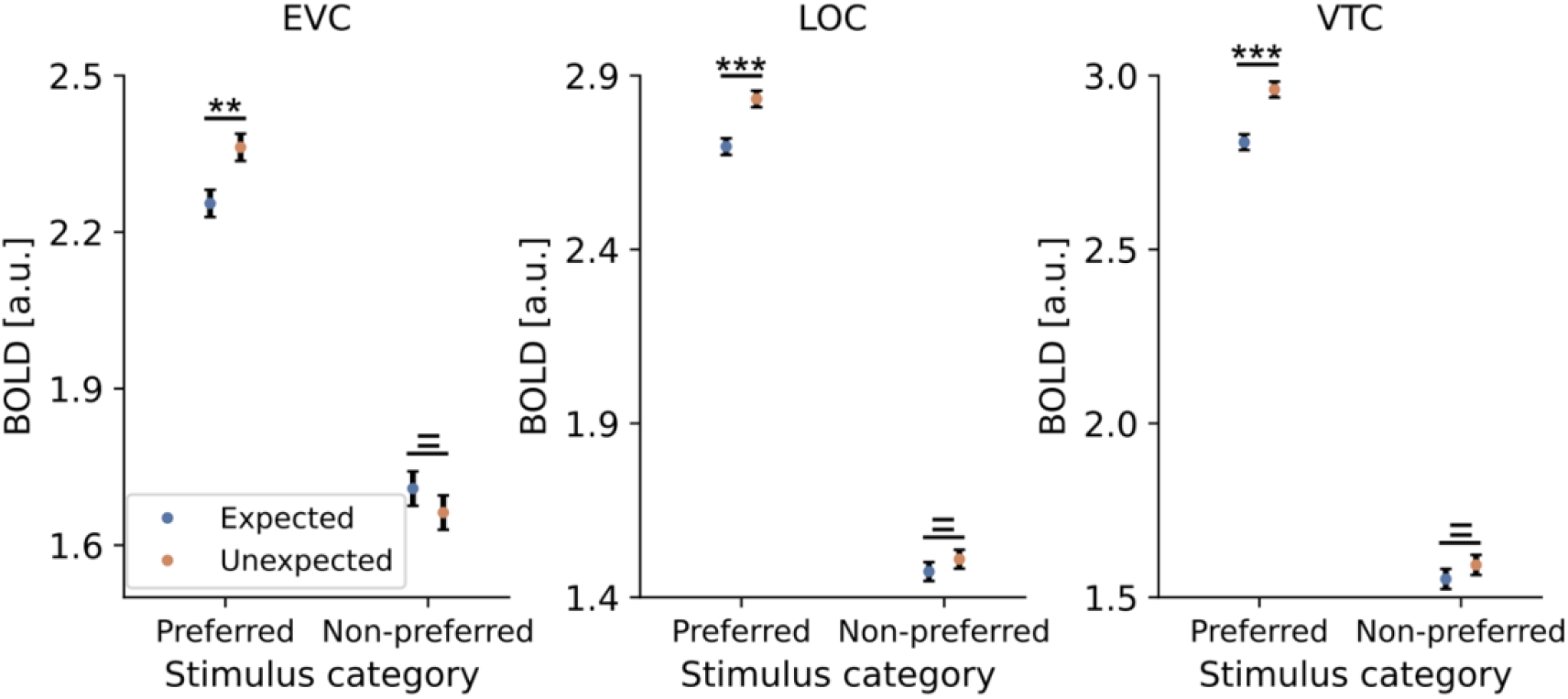
Expectation suppression only for preferred object stimuli. BOLD responses to expected (blue) and unexpected (orange) object images for preferred and non-preferred stimuli within EVC (left), LOC (middle), and VTC (right). In all three ROIs, BOLD responses were suppressed to expected object images exclusively when the object category was preferred. BOLD responses did not differ between expected and unexpected images for non-preferred object images. Error bars indicate within-subject SE. **p < 0.01, ***p < 0.001, ^=^ BF_10_ <1/3.

This pattern of results was also confirmed by a significant interaction (V1:, F_(1,33)_ = 7.507, p = 0.009, η^2^ = 0.185; LOC: F_(1,33)_ = 5.582, p = 0.024, η^2^ = 0.145; VTC: F_(1,33)_ = 7.539, p = 0.009, η^2^ = 0.186) between expectation (expected, unexpected) and preference (preferred, non-preferred). Furthermore, a Bayesian analysis showed moderate support for the absence of expectation suppression for non-preferred stimuli (V1: BF_10_ = 0.288; LOC: BF_10_ = 0.273; VTC: BF_10_ = 0.289). Crucially, neural populations showed reliable activations also to the non-preferred stimuli in all ROIs (all p < 1.0e-10, all d_z_ > 1.6; for details see Table S2), hence the absence of expectation suppression for non-preferred stimuli could not be explained by a lack of visual responsiveness. In sum, our results showed that expectations derived from conceptual associations selectively suppressed responses to preferred stimulus categories, highlighting that sensory predictions were tuning specific.

## Discussion

Sensory priors, derived from perceptual associations, play a crucial role in modulating visual processing (de Lange et al., 2018). However, less is known about how conceptual associations, for instance derived from linguistic input, influence vision. Here we examined whether conceptual associations could serve as sensory priors and how visual processing of object images is modulated by such priors. To this end, we first exposed participants to probabilistically associated word-word pairs. Participants were not informed of the underlying statistical regularities, nor required to learn them. However, the regularities could be used to facilitate performance, aiding in classification of the leading and trailing words as (in-)congruent with respect to referring to a (non-)living object. On the subsequent day, the trailing (second) words were replaced with pictures of the corresponding objects, resulting in word-image pairs.

Participants were asked to classify the object images as showing alive or not alive objects, while fMRI BOLD responses were measured. We observed faster classifications of object images corresponding to previously expected trailing words compared to unexpected ones. Crucially, during fMRI scanning object images were equiprobable, hence not allowing for any statistical learning of the word-image pairs. Thus, participants must have generalized the previously learned word-word associations to the conceptually matching word-image pairs. When analyzing the fMRI data, we observed differences in BOLD responses to object images corresponding to the previously expected trailing words compared to unexpected ones. Specifically, BOLD responses were suppressed to expected compared to unexpected objects throughout the ventral visual stream, including early visual cortex, resembling results of previous studies investigating visual statistical learning (Kaposvari et al., 2018; Richter & de Lange, 2019, for a review see: de Lange et al., 2018, but also Feuerriegel et al., 2021). This prediction induced suppression of sensory responses was stimulus specific, interacting with the tuning of the neural populations. Specifically, expectation suppression was exclusively found for preferred but not for non-preferred stimulus categories, ruling out unspecific global surprisal as source of the observed effect. Given that predictions seemed to selectively suppress neural populations tuned towards the expected stimuli, the present results are in line with previous studies showing evidence for dampened sensory representations for expected input following visual statistical learning (Kumar et al., 2017; Meyer & Olson, 2011; Richter et al., 2018). Curiously, while the behavioral expectation effect did extinguish over time, presumably reflecting the learning of the new non-predictive associations during fMRI scanning, the suppression of BOLD responses did not show clear extinction. This suggests a potential dissociation between the suppression of sensory responses and behavioral facilitation.

### Prediction errors in (early) visual cortex from conceptual category priors

Participants learned word-word pairs, hence any predictions during exposure to the word-image pairs must have originated at the level of category expectations, such as predicting the stimulus category “dogs” given a leading word. Dogs can have different shapes, colors, sizes and can assume different body positions, or more generally speaking, low-level visual features can vary dramatically yet still refer to the same object. Moreover, numerous exemplars per category were presented without exemplar repetitions within a run. Thus, it seems unlikely that category priors resulted in a predictive pre-activation of specific low-level features that are relevant for EVC representations, such as oriented edges in retinotopically specific locations (Hubel & Wiesel, 1962). Yet, we did observe tuning specific expectation suppression in EVC. How can we account for this observation?

It is possible that category expectations initially resulted in predictions of semantic and high-level visual representations of the associated object categories. That is, dogs share semantic and high-level visual features, such as being animate, having four legs, eyes and a nose. Thus, expected stimuli may result in facilitated neural processing initially by virtue of those shared features. As the predictions spread throughout the visual hierarchy (Schwiedrzik & Freiwald, 2017), EVC may subsequently be modulated by recurrent hierarchical processing. That is, if processing in higher (visual) areas is facilitated by the categorical predictions, then lower visual areas will also converge faster, or more efficiently, on a valid interpretation of the current visual stimulus due to more reliable and abundant feedback signals from higher visual areas. This explanation fits well within a hierarchical predictive coding framework (Friston, 2005; Rao & Ballard, 1999; for a review see: Walsh et al., 2020). Crucially, in the context of predictive processing theories, predictions are not necessarily about future stimuli, but are relayed top-down at each level of the cortical hierarchy, aimed at predicting bottom-up input. Hence, from this perspective, the present results may represent a faster and more efficient resolution of prediction errors throughout the visual system due to valid predictions. Specifically, conceptual and high-level visual feature predictions aid in resolving prediction errors in higher (visual) cortical areas, which through recurrent message passing across the visual hierarchy in turn also reduce prediction errors (i.e., explain away activity) in early visual cortex.

However, an explanation along the lines of hierarchical predictive coding also implies, that prediction errors are feature specific; that is representing features relevant to each level of the visual hierarchy (Clark, 2013; Friston, 2005; Walsh et al., 2020). An alternative account may hold that the observed putative prediction error signals are in fact feature unspecific modulations, representing a global gain modulation of neural responses driven by generic surprisal, following exposure to unexpected stimuli. This surprise signal, resulting in increases in arousal or attention, could then yield a proportional increase of sensory responses to the unexpected stimuli without any prediction error calculation or feature specific prediction (error) in sensory cortex (Alink & Blank, 2021). Our observation that expectation suppression was exclusively present for preferred stimulus categories contradicts this latter account, and instead favoring an explanation based on feature specific prediction errors, because an unspecific surprise signal should also be evident for non-preferred stimuli (i.e., proportional to the sensory response), which was not observed. In fact, the pattern of selective suppression for preferred stimulus categories suggest that expectations derived from conceptual associations dampen sensory representations of expected input, thus resembling previous reports of sensory dampening following visual statistical learning of object pairs (Kumar et al., 2017; Meyer & Olson, 2011; Richter et al., 2018; Richter et al., 2022).

Combined, our results add to the growing evidence, showing how predictions in the sensory brain can benefit from prior knowledge acquired from various sources (de Lange et al., 2018; Walsh et al., 2020), as in this case, priors learned in a different domain, and generalized via conceptual associations, to subsequently modulate neural processing throughout the visual hierarchy. These results also echo conclusions from previous work showing that sentences and lexical-semantic primes can modulate early sensory processing (Dikker & Pylkkanen, 2011; Hirschfeld et al., 2011).

### Sensory priors from generalized associations following incidental statistical learning

In line with the present results, prior work also showed that the expectation of higher-level attributes of complex stimuli (e.g., category expectations) can facilitate perceptual processing (Lupyan & Ward, 2013; Stein & Peelen, 2015), hence suggesting that conceptual knowledge can serve as a prior for visual processing. However, a crucial difference between previous work and the present study is that we assessed the learning, generalization, and subsequent sensory consequences of novel (arbitrary) associations following incidental statistical learning. That is, here we do not rely on well-established congruency effects, such as the word “dog” predicting the image of a dog (Gandolfo & Downing, 2019; Puri et al., 2009). Rather, we show how statistical regularities are acquired incidentally in one domain (the word “car” predicting the word “dog”) and residing at a conceptual level (i.e., abstracted from the lexical items) can subsequently serve as a sensory prior to facilitate processing of the visual features associated with the predicted image (a picture of a dog, following the word “car”). Interestingly, we observed this generalization and application of the sensory priors, even though prior work has demonstrated that statistical learning preferentially operates at less abstract (object exemplar) levels rather than generalizations across categories (Emberson & Rubinstein, 2016). Moreover, the generalization occurred in the absence of any statistical regularities during exposure to the word-image pairs. That is, the priors derived from the word-word pairs were applied to the word-image pairs even though the new statistical environment did not allow for reliable predictions, demonstrating the propensity of the brain to learn and use sensory priors. Thus, high-level conceptual associations appear to be readily used to construct concrete sensory predictions, thereby utilizing knowledge of relationships between stimuli abstracted beyond the domain in which they were initially learned.

### Distinct effects of predictions on sensory processing and behavior

Participants showed faster responses to object images that corresponded to previously expected trailing words. This behavioral facilitation was present during the initial runs and then disappeared over the course of fMRI scanning because the object images were in fact equiprobable. Thus, participants appeared to unlearn the previous associations, resulting in equal reaction times to (previously) expected and unexpected objects during the final run. Surprisingly, we did *not* find a similar extinction of the sensory suppression effect in visual cortex. Indeed, even in the final run we did find evidence for expectation suppression. Taken at face value, these results suggest that the observed sensory suppression is distinct from, and does not reflect an epiphenomenon of, behavioral facilitation. This is in line with previous reports of sensory suppression following visual statistical learning, which have been observed with task-irrelevant predictions and even when no motor responses were made (Kumar et al., 2017; Meyer & Olson, 2011; Richter et al., 2018). Moreover, it implies that the sensory priors underlying expectation suppression may differ (in their application) from those underlying the behavioral effect, and that the later decision and response stages of cortical processing may adapt to new environments before concurrent modulations in sensory cortex arise (Ferrari et al., 2022). However, we note that the neural data was noisy when analyzed on a run level, thus caution is warranted in the interpretation of these results and future work is needed to further assess the possible dissociation between the here observed behavioral and sensory effects of prediction.

### Interpretational limitations

Words referring to objects can modulate neural activity in category-selective visual areas (Kan et al., 2003; Kiefer, 2005). Moreover, natural sounds can elicit category-specific activation patterns in visual cortex (Vetter et al., 2022). These results raise the possibility that the prediction effect in the present study was induced by the leading words triggering the associated trailing *words*, which in turn may have activated corresponding category-selective visual areas. An unexpected object *image* would then lead to activation in addition to the anticipated stimulus, superficially appearing as a prediction error signal. In this case, the object associations would not need to be generalized, or operate at a conceptual level, to predict object images from the word-word pairs. Instead, processing of the object images would be modulated via the recalled word and its automatic cross-domain activation of the respective category-selective visual areas. While this account cannot be ruled out conclusively, our data speaks against this interpretation. We did not observe any expectation suppression for non-preferred stimulus categories, even though the neural populations were reliably driven by the non-preferred stimuli. If leading words automatically recall trailing words, which in turn lead to the (pre-)activation of category-selective visual areas, we would have expected to see enhanced responses to unexpected stimuli also for non-preferred stimulus categories, not selectively for preferred stimulus categories. Instead, conceptual predictions appear to selectively modulate sensory processing. In a similar vein, Vetter et al. (2022) interpreted cross-modal activations of category-specific patterns in visual cortex by naturalistic sounds in the context of predictive processing. Our data are congruent with, and can be seen as an extension of, these results by suggesting that the link between naturalistic sounds and visual cortex pattern activation may reflect a modulation via conceptual associations (i.e., sound and image referring to the same concept).

## Conclusion

In sum, our results demonstrate a widespread modulation of visual processing by conceptual associations, possibly dissociated from behavioral facilitation. Crucially, these associations were based on priors generalized across domains from words pairs to word-image pairs, following incidental statistical learning. Thus, the sensory brain appears to spontaneously use various sources for forming and applying predictions, including conceptual and high-level priors based on recently acquired conceptual associations. Such predictions subsequently modulate neural processing in a tuning specific fashion throughout the ventral visual hierarchy, including early visual cortex, resulting in a marked suppression of sensory responses to expected compared to surprising visual input.

## Materials and Methods

### Preregistration and data availability

The current study was preregistered at the Open Science Framework (OSF) before any data were analyzed. The preregistration form is available at the DOI: 10.17605/OSF.IO/RYJBN. Of the preregistration, this paper discusses how perceptual processing is influenced by statistical regularities at a conceptual level. The experiment procedures were executed as outlined in the preregistration document, unless specified otherwise in the sections below. All data and code will be shared upon publication in a peer-reviewed journal.

### Participants and data exclusion

Thirty-seven healthy participants were recruited from the Radboud University participant pool. We aimed for a sample size of 34 to obtain ≥80% power to detect a medium effect size (d = 0.5) at the standard alpha = .05. The study was approved by the local ethics committee (CMO Arnhem-Nijmegen, Radboud University Medical Center) under the umbrella ethics approval for the Donders Centre for Cognitive Neuroimaging (Imaging Human Cognition, CMO 2014/288). Written informed consent was obtained from each participant before the experiment. All subjects had normal or corrected-to-normal vision. Participants were invited for a behavioral session and an fMRI session on two consecutive days. Reimbursement was 8€/hour for the behavioral session (day 1) and 10 €/hour for the MRI session (day 2). We excluded three participants based on our pre-registered exclusion criteria. One participant was excluded from the fMRI analyses due to excessive head motion during scanning, defined as the percentage of frame-wise displacements exceeding 0.2 mm being 2 SD above the group mean. Two additional participants were excluded from all analyses due to incomplete datasets, resulting from premature termination of the experiment due to participants not complying with task instructions. The remaining 34 participants (21 female; age 25 ± 3 years, mean ± SD; range 19–34 years) were included in all analyses.

### Stimuli and experimental paradigm

#### Stimuli

We used a total of 16 objects presented as both words and images. The word stimuli subtended approximately 2–5.8 degrees of visual angle horizontally and 1 degree vertically. The length of the words was between 3 and 8 letters (5.1 ± 1.9, mean ± SD). All object images were selected by a manual search, except for the face images. Face images were chosen from the Radboud face dataset (Langner et al., 2010). The images spanned approximately 6° x 6° visual angle. All stimuli (both words and images) were from two semantic categories: living and non-living. Each category consisted of several sub-categories. Specifically, the living object words were: dog, elephant, mushroom, rose, man, woman, hand, foot. The non-living object words were: airplane, car, backpack, hat, house, church, hammer, spoon. Of these, four alive and four non-alive object words served as leading words. Similarly, four of each category were used as trailing images. For each trailing image 10 different exemplar images were used to avoid repetition of the same exemplar within a run. All stimuli were presented on a mid-gray background. Custom scripts, written in Matlab utilizing the Psychophysics Toolbox extension (Brainard, 1997; Kleiner et al., 2007; Pelli, 1997), were used for stimulus presentation. During the learning session the stimuli were presented on an LCD screen (BenQ XL2420T, 1920 × 1080 pixel resolution, 60 Hz refresh rate). During MRI scanning stimuli were presented on a MRI compatible LCD screen (Cambridge Research BOLDscreen 32, 1920 × 1080 pixel resolution, 60 Hz refresh rate), visible using an adjustable mirror mounted on the head coil.

### Procedure

Subjects participated in two sessions on two consecutive days. The first day was a behavioral session (‘learning session’) and the second day an MRI session (‘generalization session’), including an object categorization and a functional localizer task.

#### Learning session

Unbeknownst to the participants, the word-word session served as a learning session of the statistical regularities. During each trial, two words, referring to two different objects, were presented sequentially, each for 0.5 s, with an inter-trial interval (ITI) of 1-3 s randomly sampled from a uniform distribution (Figure 5A). A fixation bull’s-eye (0.3° visual angle in size) was presented throughout the experiment. The objects belonged to two groups: alive or not alive. The participants’ task was to indicate whether the two objects presented on a trial were congruent (i.e. both living or both non-living) or incongruent (i.e. one living and one non-living). Participants had to respond within 1.5 seconds after the onset of the trailing word. Feedback on behavioral performance was provided at the end of each run, using accuracy and reaction time. The button mapping was counterbalanced across participants (odd/even subject IDs). There were in total seven runs, with each run consisting of 216 trials.

**Figure 5.**
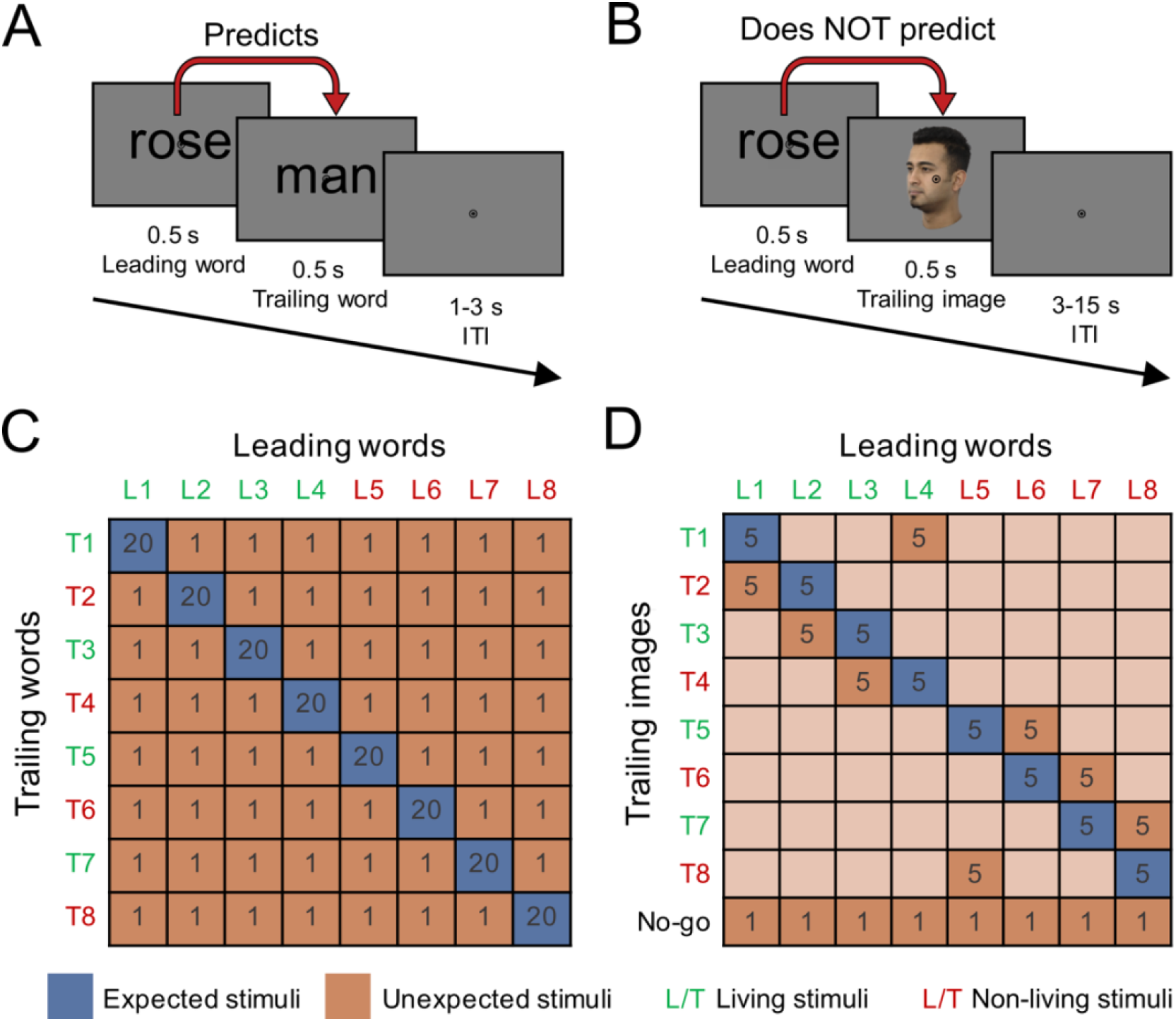
Experimental paradigm. **A)** A trial of the learning session. Two object words were presented sequentially, without ISI, each lasting 500 ms. The first word probabilistically predicted the second word. Each trial ended with a 1-3 s ITI. **B)** A trial of the generalization session. Like the learning session, two objects were presented but the trailing object word was replaced by a corresponding object image. Crucially, during this session the trailing object images were not predictable given the leading word. The leading word and trailing images were presented sequentially for 500 ms each, without ISI, followed by a 3-15 s ITI. **C)** The transitional probability matrix of the learning session, determining the associations between word pairs. L1 to L8 represent leading words and T1 to T8 represent the trailing words. Green labels indicate that the word refers to a living object while red indicates a non-living object. Blue and orange cells denote expected and unexpected word pairs respectively. The number inside each cell indicates the number of trials in the corresponding conditions per run. **D)** The transitional probability matrix of the generalization session. The matrix was identical to the learning session except for 3 changes. First, T1-T8 represent trailing images instead of words. Moreover, a no-go condition was added in which the trailing images were of the same object as the leading words. Finally, the leading words were no longer predictive of the specific trailing stimulus, instead one of two equiprobable trailing images were associated with each leading word, one of which was the previously expected object category. Thus, the blue cells represent the object images corresponding to the (previously) expected words, while the orange cells represent object images that correspond to the unexpected words.

Expectations were induced by manipulating the transitional probabilities of the word pairs. The transitional probabilities and the object words themselves were determined by an 8 × 8 transitional probability matrix (Figure 5C). For each leading word, the paired (expected) trailing word occurred with a probability of ∼74% while any other trailing word (unexpected) appeared with a probability of ∼3.7%. The specific pairings of leading and trailing words were randomized for each participant. Eight expected word pairs were randomly selected for each subject and balanced with respect to button mappings, i.e., half of the object pairs were of the same category (both alive, or both not alive) while the other half was from different categories.

On the next day, the participants performed an additional learning session in order to consolidate the learned associations. This session included two runs of the word-word task outside of the MRI scanner and another 104 trials inside the MRI scanner, during which an anatomical image was acquired.

#### Generalization (MRI) session

During fMRI scanning participants were exposed to word-image pairs. The leading words were the same as in the learning session, while the trailing words were replaced by corresponding object images. Each stimulus was presented for 500 ms followed by a variable ITI of 3–15 seconds (mean = 5 s), randomly sampled from a truncated exponential distribution. Participants indicated whether the trailing object was alive or not alive by button press. Additionally, we added a no-go condition, requiring participants to withhold their responses when the leading word and trailing image referred to the same object (e.g. the word “dog” followed by an image of a dog). We also modified the transitional probabilities. Specifically, expected pairs remained the same, but one unexpected trailing image was selected for each leading word. Crucially, the expected and unexpected trailing images were now equiprobable, appearing equally often given a leading word. Therefore, the status of a trailing object image as expected or unexpected depended completely on the participants, spontaneously and without instruction, generalizing the associations learned during the learning session to the word-image pairs. For each trailing object category we used 10 distinct exemplars, with each exemplar being shown only once as expected and once as unexpected image per run. Moreover, the exemplars varied in orientation, color, shape, and other low-level visual features to reduce the possibility that participants associated specific low-level visual features with the object category. The selected expected and unexpected trailing images were always from different categories with respect to being (not) alive, such that participants could not predict the button response based on the leading words. Participants had to respond within 1.5 seconds after the onset of the trailing images. Feedback on accuracy and reaction time were provided at the end of each run. Participants first performed a brief practice run (12 trials, ∼2 min), which was followed by four runs of the main task. Each run (∼9 min) consisted of 88 trials, including 40 trials per expectation condition and 8 no-go trials. The presentation order was fully randomized.

#### Functional localizer

Finally, participants performed a functional localizer consisting of three types of stimuli: words, intact images, and globally phase-scrambled images. Participants were instructed to respond by button press whenever two consecutive images were identical while keeping their gaze at the central fixation point. The task consisted of three runs in a block design. Each run (∼8 min) included 16 word blocks, 16 intact image blocks, 8 phase-scrambled image blocks, and 4 null-event blocks. Each stimulus type was presented fifteen times per block, each time flashing at 1.4 Hz (500 ms on, 200 ms off) for 10.5 s. The order of blocks (intact and phase-scrambled images) was fully randomized.

#### fMRI data acquisition

Functional and anatomical images were collected on a 3T Skyra MRI system (Siemens), using a 32-channel head coil. The functional images were acquired using a whole-brain T2*-weighted multiband-4 sequence (TR/TE = 1500/33.4 ms, 68 slices, voxel size 2 mm isotropic, FOV = 210 mm, 75° flip angle, A/P phase encoding direction, bandwidth = 2090 Hz/Px). The anatomical images were acquired with a T1-weighted MP-RAGE (GRAPPA acceleration factor = 2, TR/TE = 2300/3.03 ms, voxel size 1 mm isotropic, 8° flip angle).

### Data analysis

#### Behavioral data analysis

Behavioral data were analyzed in terms of accuracy and RTs. The expectation benefits were defined as higher accuracy (expected – unexpected) and faster RTs (unexpected – expected) to expected stimuli. The accuracy was calculated separately for expected and unexpected trials for each subject and contrasted with a paired t-test. For RT analysis, only correct responses were analyzed. The trials with RTs faster than 100 ms and slower than 1500 ms were rejected as outliers from further analysis. These RTs were then averaged for each expectation condition separately per participant and subjected to a paired t-test across subjects. The effect size of the difference was calculated in terms of Cohen’s d_z_ (Lakens, 2013). A Bayesian t-test with a Cauchy prior width of 0.707 was used to assess any not statistically significant results to assess evidence for absence of an effect. All standard errors of the mean (SEM) were calculated as the within-subject normalized SEM (Cousineau, 2005). Since the unexpected trailing word trials can required a change in the response during the learning session, we additionally assess expectation effects without response adjustments. To this end we split the unexpected trailing word trials into two groups depending on whether they required the same button response as the expected trailing words or not.

#### fMRI data preprocessing

fMRI data preprocessing was performed using FSL 5.0.9 (FMRIB Software Library; Oxford, UK; www.fmrib.ox.ac.uk/fsl; RRID:SCR_002823). The preprocessing pipeline included brain extraction (BET), motion correction (MCFLIRT) and temporal high-pass filtering (128 s). For the univariate analysis, the data were spatial smoothed with a Gaussian kernel (5 mm FWHM). For the multivariate analysis, no spatial smoothing was applied. Functional images were registered to the anatomical image using boundary-based registration (BBR) as implemented in FLIRT and subsequently normalized to the MNI152 T1 2 mm template brain using linear registration with 12 degrees of freedom. To allow for signal stabilization, the first four volumes of each run were discarded.

#### Univariate data analysis

We modeled BOLD signal responses to the different experimental conditions by fitting voxel-wise GLMs to each run’s data, using FSL FEAT. We modeled the events of interest (expected, unexpected, and no-go) as three separate regressors with a duration of one second (the combined duration of leading word and trailing image) and convolved them with a double gamma hemodynamic response function. The contrast of interest was expected minus unexpected images, hence negative parameter estimates indicating expectation suppression. In addition, the first-order temporal derivatives of the regressors of interest and 24 motion regressors (FSL’s standard + extended set of motion parameters) were added as nuisance regressors. FSL’s fixed effects analysis was used to combine data across runs while the FSL’s mixed effects analysis (FLAME 1) was used to combine data across participants. Gaussian random-field cluster thresholding was used to correct for multiple comparisons, using the default settings of FSL, with a cluster formation threshold of p < 0.001 (z ≥ 3.1) and a cluster significance threshold of p < 0.05.

#### Region of interest (ROI) analysis

Three ROIs (EVC, LOC and VTC) were selected to examine expectation suppression in the ventral visual stream. For each ROI the mean parameter estimates were extracted separately for expected and unexpected conditions per participant in native space (i.e., without normalization to MNI space). The parameter estimates were then divided by 100 to yield percent signal change relative to baseline (Mumford, 2007). These mean parameter estimates were submitted to paired t-tests for each ROI. Cohen’s d_z_ was calculated as a measure of effect size. Additionally, we performed the ROI analyses for each run separately to assess the expectation effect over time.

#### ROI definition

To examine the expectation effects throughout the visual hierarchy, we defined three ROIs: EVC, object-selective LOC, and category-selective VTC. The preregistered EVC and LOC ROIs were used to investigate modulations by expectation at the low-level and intermediate object-selective visual regions. In addition to the ROIs from the preregistration, we added VTC to further examine potential activity modulations by expectation in higher category-selective regions that are sensitive to the distinction between living and non-living objects (Thorat et al., 2019).

EVC was defined as the union of V1 and V2. Freesurfer 6.0 (General Hospital Corporation, Boston MA, USA, RRID: SCR_001847) was used to extract labels for V1 and V2 per subject based on their anatomical image. These labels were then transformed to native EPI space and combined into a bilateral mask. Object selective LOC was defined as voxels within an anatomical LOC mask, derived from the Harvard-Oxford cortical atlas, that were more responsive to intact compared to phase-scrambled objects. To this end, we modeled the three types of stimuli during the fMRI localizer runs (words, intact and phase-scrambled objects) with their corresponding duration (10.5 s). First-order temporal derivatives and 24 motion regressors were added as nuisance regressors. The contrast of the intact objects minus scrambled objects was thresholded at z >= 4.3 (one-sided, p < 1e-5) and further constrained by the anatomical LOC mask. The resulting LOC masks all contained at least 600 voxels in native space per participant. The VTC ROI mask was created using anatomical masks from the Harvard-Oxford cortical atlas, including the temporal-occipital fusiform cortex, the temporal gyrus, and the parahippocampal gyrus. The resulting mask was transformed from MNI space to the participants native space using FSL FLIRT. Finally, we constrained each of the ROI masks to the most informative voxels regarding object image category. Specifically, we decoded object category during the functional localizer run per participant (see: Multi-voxel pattern analysis), and then for each ROI and subject selected the 300 voxels forming the most informative neighborhood in the decoding analysis. As a robustness check for all ROI analyses, we repeated each analysis with mask sizes ranging from 50 to 600 voxels in steps of 50 voxels.

#### Expectation suppression selectivity analysis

In addition to the main expectation analysis, we also performed an analysis assessing the selectivity of the expectation modulation. First, a GLM based analysis was performed using the data from the generalization session. However, now we modeled the expected and unexpected events for each aliveness category separately, thus resulting in four regressors of interest (expected alive, unexpected alive, expected not alive and unexpected not alive). Additionally, we also modeled the no-go condition and the nuisance regressors (same as for other analyses). Next, for the three anatomical masks (EVC, LOC, and VTC) we selected voxels that preferentially responded to images of living or non-living objects. Preference was estimated using the independent localizer data, for which we modeled the image stimuli separately for living and non-living objects. Then the contrast “living – non-living” was used to identify 300 voxels that were more responsive (highest z-scores) to alive compared to not alive objects. The inverse contrast (non-living – living) was used to identify 300 voxels that were more responsive to images of non-living objects. The ROI masks, split into living preferred and non-living preferred, were then applied to the main task data to obtain the mean parameter estimates for expected and unexpected trailing images per aliveness category. The mean parameter estimates were then averaged by stimulus preference. In other words, living stimuli contributed to living preferring voxels together with non-living stimuli for non-living preferring voxels. Thus, this analysis results in four datapoints per participant and ROI (EVC, LOC, VTC): expected preferred stimuli, unexpected preferred stimuli, expected non-preferred stimuli and unexpected non-preferred stimuli. For each ROI, the data were submitted to a 2 × 2 repeated measures ANOVA with expectation (expected, unexpected) and preference (preference or non-preference) as factors. The BOLD response difference between expected and unexpected trials for each preference category was examined using simple paired t-tests. Partial eta-squared (η^2^) and Cohen’s d_z_ were calculated as effect size for the RM ANOVA and paired t-test respectively. Bayesian t-tests were used to assess evidence for the absence of an effect.

#### Multi-voxel pattern analysis

For the multi-voxel pattern analyses, no spatial smoothing was applied. Parameter estimate maps per localizer trial were estimated using a GLM based LS-S approach (Mumford et al., 2012). Each model contained four regressors of interest: one for the trial of interest and three for all other trials per condition (expected, unexpected and no-go). The resulting parameter estimate maps were subsequently used in a searchlight analysis with a sphere of 6 mm radius. A leave-one-run-out classifier, using linear SVM, was fit to the four object categories (i.e. faces, body parts, buildings, and tools) as classes. The classifier was trained and tested on the independent localizer data across the whole brain for each participant in native space. The resulting decoding accuracy maps were then used to constrain the ROI masks per participant.

## Software

PsychToolbox (Brainard, 1997; Kleiner et al., 2007; Pelli, 1997) running on MATLAB R2017b (The MathWorks, Natick, MA, USA, RRID:SCR_001622) was used for stimuli presentation. (f)MRI data preprocessing and analysis was performed using FSL 5.0.9 (FMRIB Software Library; Oxford, UK; RRID:SCR_002823) and Freesurfer 6.0 (General Hospital Corporation, Boston MA, USA, RRID:SCR_001847). Python 3.7.4 (Python Software Foundation, RRID:SCR_008394) was used for data processing with following libraries: NumPy 1.17.2 (van der Walt et al., 2011), Pandas 0.25.1 (McKinney, 2010), nilearn 0.8.1 (Abraham et al., 2014), Scikit-learn 0.18.1 (RRID:SCR_002577); Matplotlib 3.1.1 (Brett et al., 2019) and Nanslice (Hunter, 2007) were used for data visualization. Pingouin 0.2.9 (Wood, 2017/2020) was used for statistical tests, including RM ANOVA, paired t-tests and Bayesian analyses.

## Author contributions

C.Y., D.R. and F.P.d.L. designed research; C.Y. performed experiments; C.Y. and D.R. analyzed data; C.Y. and D.R. wrote the first draft of the paper; C.Y., D.R. and F.P.d.L. edited and revised the paper.

## Acknowledgements

We thank Xueyan Jiang for assistance with data acquisition. This study was supported by the EC Horizon 2020 Program ERC Starting Grant 678286 “Contextvision” awarded to F.P.d.L. and China Scholarship Council CSC201708330238 awarded to C.Y.

## Supplementary Information

**Figure S1.**
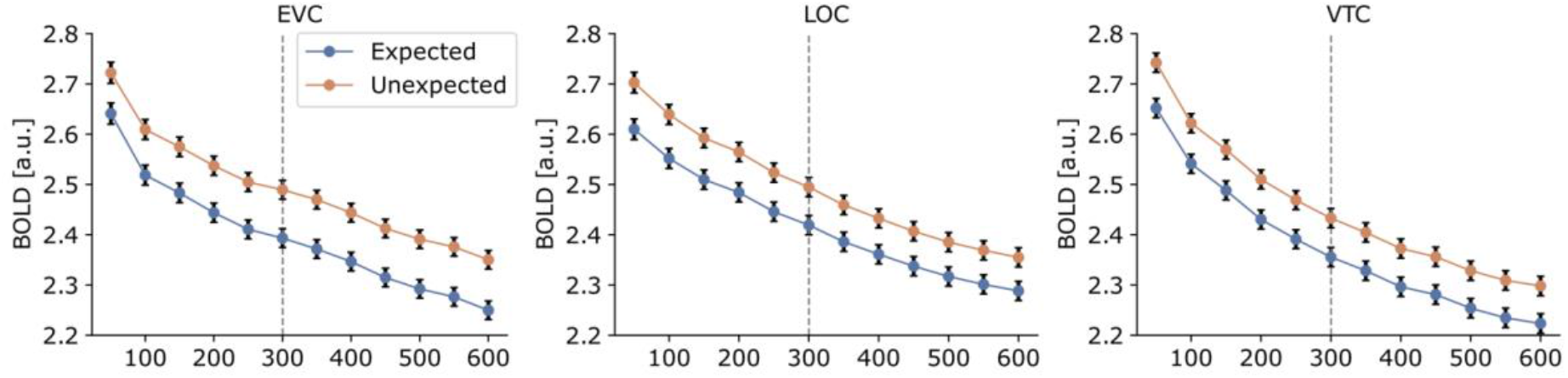
Expectation suppression over a range of ROI sizes. Averaged BOLD responses (parameter estimates) to expected (blue) and unexpected (orange) object images within EVC, LOC, and VTC for varying ROI sizes from 50 to 600 voxels. Vertical dashed lines indicate the pre-registered ROI size of 300 voxels, as used in Figure 2C.

**Table S1.** Expectation suppression over runs. Results of paired t-tests comparing the BOLD responses to expected and unexpected object images for each run in all three ROIs.

|  | Run 1 | Run 2 | Run 3 | Run 4 |
| --- | --- | --- | --- | --- |
| EVC : | $t_{(33)} = 1.264$,<br>$p = 0.215$,<br>$d_z = 0.217$ | $t_{(33)} = 2.514$,<br>$p = 0.017$,<br>$d_z = 0.431$ | $t_{(33)} = 0.528$,<br>$p = 0.601$,<br>$d_z = 0.091$ | $t_{(33)} = 2.006$,<br>$p = 0.053$,<br>$d_z = 0.344$ |
| LOC: | $t_{(33)} = 0.435$,<br>$p = 0.666$,<br>$d_z = 0.075$ | $t_{(33)} = 2.067$,<br>$p = 0.047$,<br>$d_z = 0.355$ | $t_{(33)} = 0.067$,<br>$p = 0.947$,<br>$d_z = 0.012$ | $t_{(33)} = 2.227$,<br>$p = 0.033$,<br>$d_z = 0.382$ |
| VTC: | $t_{(33)} = 0.313$,<br>$p = 0.756$,<br>$d_z = 0.054$ | $t_{(33)} = 2.584$,<br>$p = 0.014$,<br>$d_z = 0.443$ | $t_{(33)} = 0.153$,<br>$p = 0.879$,<br>$d_z = 0.026$ | $t_{(33)} = 2.095$,<br>$p = 0.044$,<br>$d_z = 0.359$ |

**Table S2.** Robust neural activation to non-preferred stimuli. Results of one-sample tests comparing the BOLD responses to non-preferred stimuli against no visual stimulation for each expectation condition in all ROIs. Results demonstrate that voxels in all three ROIs significantly responded to non-preferred stimulus categories.

|  | Expected | Unexpected |
| --- | --- | --- |
| EVC: | $t_{(33)} = 10.773$, $p = 2.4e-12$, $d_z = 1.847$ | $t_{(33)} = 9.594$, $p = 4.5e-11$, $d_z = 1.645$ |
| LOC: | $t_{(33)} = 11.059$, $p = 1.2e-12$, $d_z = 1.897$ | $t_{(33)} = 12.403$, $p = 5.7e-14$, $d_z = 2.127$ |
| VTC: | $t_{(33)} = 12.954$, $p = 1.7e-14$, $d_z = 2.222$ | $t_{(33)} = 14.512$, $p = 7.1e-16$, $d_z = 2.487$ |

**Table S3.** Reliable decoding of stimulus category in all visual ROIs. Results of one-sample t-tests comparing the decoding accuracies against chance level (50%) for each expectation condition in all ROIs. Results show that object category could be decoded reliably within all ROIs.

|  | Expected | Unexpected |
| --- | --- | --- |
| EVC: | $t_{(33)} = 11.917, p = 1.7e-13, d_z = 2.044$ | $t_{(33)} = 15.206, p = 1.8e-16, d_z = 2.608$ |
| LOC: | $t_{(33)} = 15.560, p = 9.3e-17, d_z = 2.669$ | $t_{(33)} = 14.256, p = 1.2e-15, d_z = 2.445$ |
| VTC: | $t_{(33)} = 19.823, p = 6.7e-20, d_z = 3.400$ | $t_{(33)} = 16.318, p = 2.3e-17, d_z = 2.799$ |

## Notes

### Competing Interest Statement

The authors have declared no competing interest.

